# TMEM120A/TACAN inhibits mechanically activated Piezo2 channels

**DOI:** 10.1101/2021.06.30.450616

**Authors:** John Smith Del Rosario, Matthew Gabrielle, Yevgen Yudin, Tibor Rohacs

## Abstract

Mechanically activated Piezo2 channels are key mediators of light touch and proprioception in mice and humans. Relatively little is known about what other proteins regulate Piezo2 activity in a cellular context. TACAN (TMEM120A) was proposed to act as a high threshold mechanically activated ion channel in nociceptive dorsal root ganglion (DRG) neurons. Here we find that TACAN co-expression robustly reduced mechanically activated Piezo2 currents, but did not inhibit mechanically activated Piezo1 and TREK1 currents. TACAN co-expression did not affect cell surface expression of either Piezo1 or Piezo2 and did not have major effects on the cortical actin or tubulin cytoskeleton. TACAN expression alone did not result in the appearance of mechanically activated currents above background. In addition, TACAN and Piezo2 expression in DRG neurons overlapped, and siRNA mediated knockdown of TACAN did not decrease the proportion of slowly adapting mechanically activated currents, but resulted in an increased proportion of rapidly adapting currents. Our data do not support TACAN being a mechanically activated ion channel, and identify it as a negative modulator of Piezo2 channel activity.

## INTRODUCTION

Piezo2 is a non-selective cation channel that is responsible for the rapidly adapting mechanically activated currents in DRG neurons (Coste et al., 2010). Piezo2 plays crucial roles in light touch and proprioception both in mice and in humans (Kefauver et al., 2020). In humans, loss of function mutations in Piezo2 lead to impaired proprioception, loss of discriminative touch perception, as well as ataxia and muscular dystrophy (Chesler et al., 2016). In mice, combined deletion of Piezo2 in DRG neurons and in Merkel cells, resulted in a profound loss of light touch (Ranade et al., 2014). Selective deletion of Piezo2 in proprioceptive neurons in mice recapitulated not only ataxia, but also much of the skeletal abnormalities observed in human loss of function patients (Assaraf et al., 2020).

In contrast to its well-established roles in light touch, and proprioception, the role of Piezo2 channels in detecting painful stimuli is less clear. Deletion of Piezo2 in DRG neurons and Merkel cells resulted in no change in sensitivity to noxious mechanical stimuli and mechanical allodynia in inflammation (Ranade et al., 2014). Deletion of Piezo2 in a different subset of DRG neurons, on the other hand, reduced sensitivity to noxious mechanical stimuli, as well as reduced inflammatory and neuropathic mechanical allodynia (Murthy et al., 2018b). In another study deletion of Piezo2 in DRG neurons increased threshold to light touch, but paradoxically reduced the threshold to painful mechanical stimuli, indicating that activation of Piezo2 channels may reduce pain (Zhang et al., 2019), in accordance with the gate control theory of pain (Melzack and Wall, 1965). These data highlight the potentially complex role of Piezo2 in detecting noxious mechanical stimuli.

Piezo2 is not the only mechanically activated channel in DRG neurons, and it is not the sole sensor responsible for all aspects of somatosensory touch and mechanical pain (Kefauver et al., 2020). Several novel putative mechanically activated channels have been identified in recent years, including Tentonin3 (TMEM150C) (Hong et al., 2016), Elkin1 (TMEM87A) (Patkunarajah et al., 2020), OSCAs (TMEM63) (Murthy et al., 2018a) and TACAN (TMEM120A) (Beaulieu-Laroche et al., 2020). The physiological roles of these novel putative mechanically activated channels is not very well understood.

Tentonin3 (TMEM150C) was proposed to be a component of a slowly adapting mechanically activated currents in proprioceptive DRG neurons, and its genetic deletion resulted in abnormal gait and loss of motor coordination (Hong et al., 2016). Later its function as a mechanically activated channel was debated (Dubin et al., 2017; Hong et al., 2017) and it was proposed that Tentonin3 acts as a modulator of Piezo2 and Piezo1 channel activity (Anderson et al., 2018).

TACAN (TMEM120A) was also proposed to be responsible for the slowly adapting mechanically activated currents in DRG neurons (Beaulieu-Laroche et al., 2020). Cells transfected with TACAN displayed mechanically activated currents above background when stimulated by negative pressure in the cell attached mode, but not when stimulated by indentation with a glass probe in whole cell patch clamp experiments (Beaulieu-Laroche et al., 2020). Inducible genetic deletion of TACAN in non-peptidergic Mrgprd positive DRG neurons in mice resulted in reduced behavioral responses to painful mechanical stimuli (Beaulieu-Laroche et al., 2020). TACAN knockdown in DRG neurons by intrathecal administration of antisense oligodeoxynucleotides in rats reduced mechanical hyperalgesia induced by intradermal administration of various proinflammatory compounds, but had no effect on mechanical allodynia in chemotherapy-induced neuropathic pain (Bonet et al., 2020).

Here we examined whether TACAN can act as a modulator of Piezo channels. We found that coexpression of TACAN with Piezo2 channels resulted in a robust reduction in mechanically activated currents in whole cell patch clamp experiments compared to cells expressing Piezo2 alone. On the other hand, coexpressing TACAN with Piezo1 did not have a significant effect on mechanically induced currents either when the cells were stimulated with a blunt glass probe in the whole cell configuration, or when negative pressure was used in cell attached patches. Expressing TACAN alone did not induce mechanically activated currents above background levels observed in control cells. Co-expressing TACAN with Piezo1 or Piezo2 channels, did not alter their cell surface localization, indicating that TACAN altered the gating of Piezo2 not its abundance in the membrane. TACAN expression in DRG neurons overlapped with Piezo2, and siRNA mediated knockdown of TACAN did not decrease the proportion of slowly adapting currents, but it increased the proportion of rapidly adapting mechanically activated currents in DRG neurons. Our data identify TACAN as a negative modulator of Piezo2 channels.

## RESULTS

We first transiently transfected HEK293 cells with Piezo2 and TACAN, and measured mechanically activated currents in response to indentation of the cell membrane with a blunt glass probe in whole cell patch clamp experiments. Cells expressing Piezo2 alone displayed rapidly adapting mechanically activated currents that showed increased amplitudes in response to deeper indentations (**Fig. 1A-C**). When TACAN was co-transfected with Piezo2, the majority of the cells did not show responses to mechanical indentations, with a small number of cells responding to stronger stimuli **(Fig. 1A-C)**. To ensure that all cells we patched in the TACAN transfected cells indeed expressed TACAN and Piezo2, we transfected HEK293 cells with TACAN tagged with tdTomato, and Piezo2 tagged with GFP, and patched cells displaying both red and green fluorescence. Coexpression of tdTomato-TACAN robustly suppressed Piezo2 currents **(Fig. 1D-F)** indicating that the responses in the TACAN transfected group were unlikely to be due to the lack of TACAN expression, and the lack of responses were not due to the lack of Piezo2. To ensure that the reduction in Piezo2 channel activity is not specific to HEK293 cells, we also co-expressed TACAN with Piezo2 in Neuro2A (N2A) cells in which Piezo1 was deleted with CRISPR (Moroni et al., 2018; Romero et al., 2020). In these cells we could perform deeper indentations of the cell without losing the seal, thus we detected larger currents. Similar to HEK293 cells, co-expression of tdTomato-TACAN with GFP-Piezo2 strongly reduced mechanically activated currents **(Fig. 1G-I)**.

**Figure 1:**
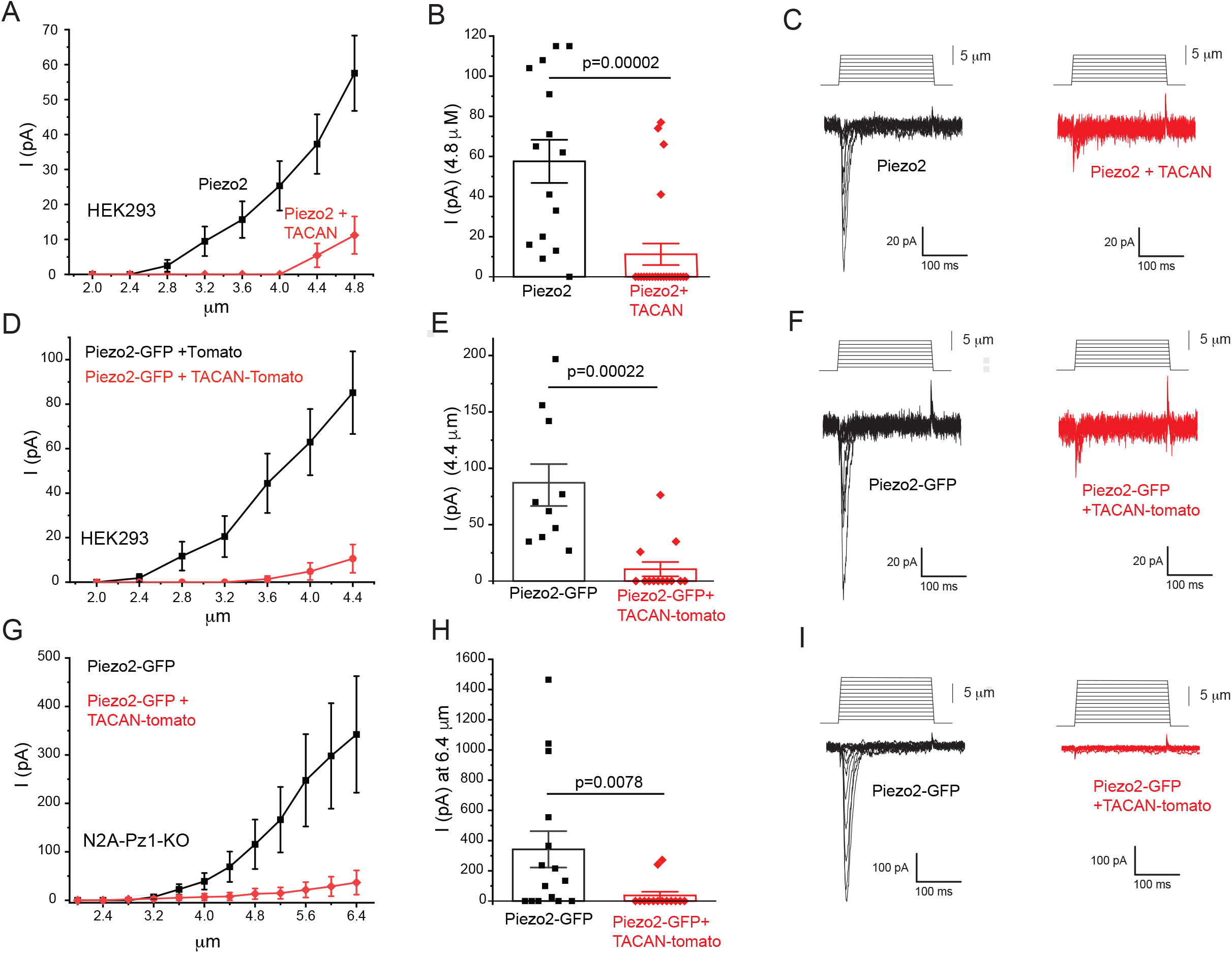
TACAN inhibits Piezo2 currents. Whole cell patch clamp experiments at -60 mV in cells expressing Piezo2 with, or without TACAN were performed as described in the methods section. **(A)** HEK293 cells were transfected with Piezo2 and GFP with or without TACAN. In some cells GFP-tagged Piezo2 was used instead of Piezo2 plus GFP. Current amplitudes are plotted (mean ± SEM) for Piezo2 expressing cells (black) and for cells expressing Piezo2 and TACAN (red). **(B)** Scatter plots and mean ± SEM for current amplitudes at 4.8 μm indentation. Statistical significance was calculated with the Mann Whitney test. **(C)** Representative current traces. **(D)** HEK293 cells were transfected with GFP-tagged Piezo2 and with tdTomato-tagged TACAN or tdTomato. Current amplitudes are plotted (mean ± SEM) for Piezo2 expressing cells (black) and for cells expressing Piezo2 and TACAN (red). **(E)** Scatter plots and mean ± SEM for current amplitudes at 4.4 μm indentation. Statistical significance was calculated with the Mann Whitney test. **(F)** Representative current traces. **(G)** Piezo1 deficient Neuro2A cells were transfected with GFP-tagged Piezo2 and with tdTomato-tagged TACAN or tdTomato. Current amplitudes (mean ± SEM) are plotted for Piezo2 expressing cells (black) and for cells expressing Piezo2 and TACAN (red). **(H)** Scatter plots and mean ± SEM for current amplitudes at 6 μm indentation. Statistical significance was calculated with the Mann Whitney test. **(I)** Representative current traces.

We also patched HEK293 cells expressing tdTomato-TACAN alone, and we did not observe any mechanically activated currents in response to indentation with a blunt glass probe up to 9.2 μm (data not shown, n=11), in accordance with the original report describing TACAN (Beaulieu-Laroche et al., 2020).

Both in HEK293 cells and in N2A cells we stimulated all cells with increasing indentations until the seal was lost, but only plotted data in Figure 1 up to the indentation depth where all cells still had intact seals. **Figure 1 Figure Supplement 1** displays the full data showing the overall much lower responsiveness of TACAN expressing cells and representative traces for mechanically activated currents in TACAN expressing cells at higher indentation levels. Piezo2 currents in cells expressing either Piezo2-GFP alone or with tomato-tagged TACAN showed similar inactivation kinetics in both HEK293 cells and in N2A cells **(Figure 1 Figure Supplement 1H,L)**. In HEK293 cells co-transfected with Piezo2 and non-Tagged TACAN, the time constant of inactivation increased slightly but significantly **(Figure 1 Figure Supplement 1D)**.

We also tested if the TACAN homologue TMEM120B has any effect on Piezo2 current amplitudes. **Figure 1 Figure Supplement 1M,N** shows that Piezo2 current amplitudes were similar in HEK293 cells transfected with or without TMEM120B.

Next, we tested if TACAN modulates the closely related Piezo1 channels. Co-expression of TACAN in HEK293 cells did not inhibit Piezo1 channel activity evoked by indentation with a blunt glass probe in whole cell patch clamp experiments (**Fig. 2A-C, Figure 2 Figure Supplement 1A,B**.).

**Figure 2:**
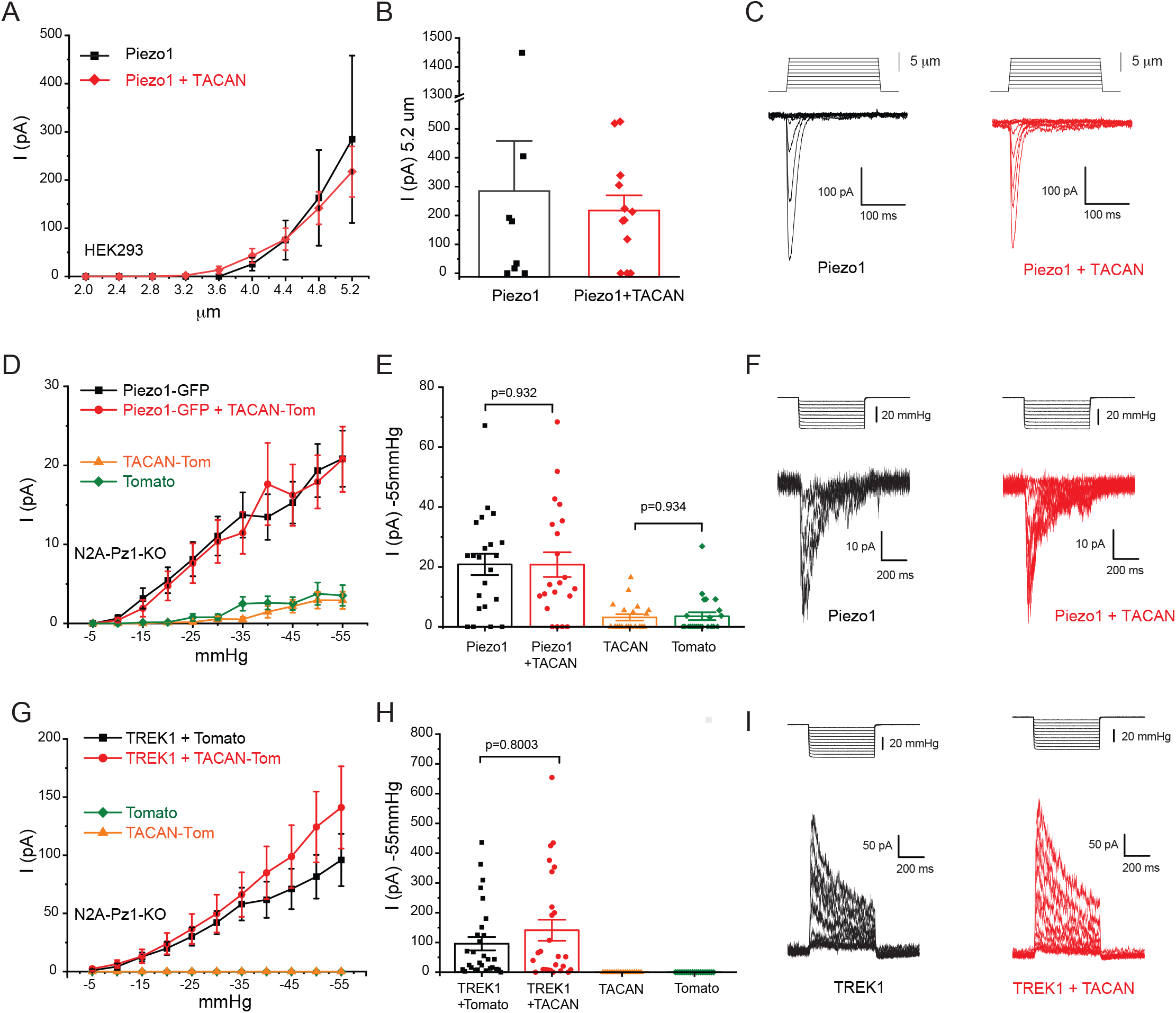
TACAN does not inhibit Piezo1 and TREK1 currents. **(A)** HEK293 cells were transfected with Piezo1 in IRES-GFP vector, with or without TACAN. Mechanically activated currents were evoked by increasing indentations with a blunt glass probe in whole cell patch clamp experiments. Current amplitudes are plotted (mean ± SEM) for Piezo1 expressing cells (black) and for cells expressing Piezo1and TACAN (red). **(B)** Scatter plots and mean ± SEM for current amplitudes at 5.2 μm indentation. Statistical significance was calculated with the Mann Whitney test. **(C)** Representative current traces. **(D)** Piezo1 deficient Neuro2A cells were transfected with GFP-Piezo1 with or without tdTomato-TACAN, and with tdTomato-TACAN alone or tdTomato alone. Mechanically activated currents were evoked by applying increasing negative pressures through the patch pipette in cell-attached patch clamp experiments. Measurements were performed at -80 mV holding potential. Current amplitudes are plotted (mean ± SEM) for cells expressing Piezo1 (black), Piezo1 and TACAN (red), TACAN alone (orange) and tdTomato alone (green). **(E)** Scatter plots and mean ± SEM for current amplitudes at -55 mmHg. Statistical significance was calculated with the Mann Whitney test. **(F)** Representative current traces. **(G)** Piezo1 deficient Neuro2A cells were transfected with TREK1 with, or without tdTomato-TACAN, and with tdTomato-TACAN alone or tdTomato alone. Mechanically activated currents were evoked by applying increasing negative pressures through the patch pipette in cell attached patch clamp experiments. Measurements were performed at 0 mV holding potential. Current amplitudes are plotted (mean ± SEM) for cells expressing TREK1 (black), TREK1 and TACAN (red), TACAN alone (orange) and tdTomato alone (green). **(H)** Scatter plots and mean ± SEM for current amplitudes at -55 mmHg. Statistical significance was calculated with the Mann Whitney test. **(I)** Representative current traces.

Unlike Piezo2 (Shin et al., 2019), Piezo1 currents can be reliably evoked by negative pressure in the cell attached mode (Coste et al., 2010). To test the effect of TACAN in this modality, we expressed GFP-Piezo1 with tdTomato-TACAN or tdTomato in Piezo1 deficient Neuro2A cells. Negative pressures applied through the patch pipette reproducibly evoked mechanically activated currents. Current amplitudes in the TACAN expressing cells were similar to those without TACAN (**Fig. 2D-F**). We also performed experiments in cells only expressing TACAN, or tdTomato. In contrast to Piezo1 expressing cells, both groups displayed no, or negligible currents at low pressures. Increasing negative pressures evoked small currents in both tdTomato-TACAN and in tdTomato transfected cells, but the amplitudes of those were very similar in the two groups **(Fig. 2D,E, Figure 2 Figure Supplement 1E-H)**.

We also tested if TACAN had any effect on the activity of TREK1, a mechanically activated K^+^ selective ion channel. We evoked outward TREK1 currents in cell attached patches in N2A cells by negative pressures at 0 mV at physiological extracellular and intracellular K^+^ concentrations (**Fig. 2G-I**). Currents in cells cotransfected with tdTomato-TACAN and TREK1 showed similar current amplitudes to those transfected with TREK1 and tdTomato **(Fig. 2G-I, Figure 2 Figure Supplement 1I-L)**. In cells expressing tdTomato-TACAN, or tdTomato alone, negative pressures did not induce any currents in these conditions **(Fig. 2G,H, Figure 2 Figure Supplement 1K,L)**.

Next, we tested if the inhibition of Piezo2 currents by TACAN was caused by decreased cell surface expression. For this, we cotransfected HEK293 cells with GFP-tagged Piezo channels, and tdTomato-TACAN or tdTomato. To assess cell surface expression, we performed dual color Total Internal Reflection Fluorescence (TIRF) imaging. In TIRF imaging only a narrow layer at the bottom of the cell is illuminated, representing the plasma membrane and a narrow sub plasma membrane region. **Figure 3A,B** shows that TIRF intensity of GFP-Piezo2 was similar in cells cotransfected with tdTomato TACAN or tdTomato, indicating that coexpression of TACAN did not change the cell surface expression of Piezo2. **Figure 3D,E** shows that coexpression of tdTomato-TACAN did not change the cell surface expression of GFP-Piezo1 either.

**Figure 3:**
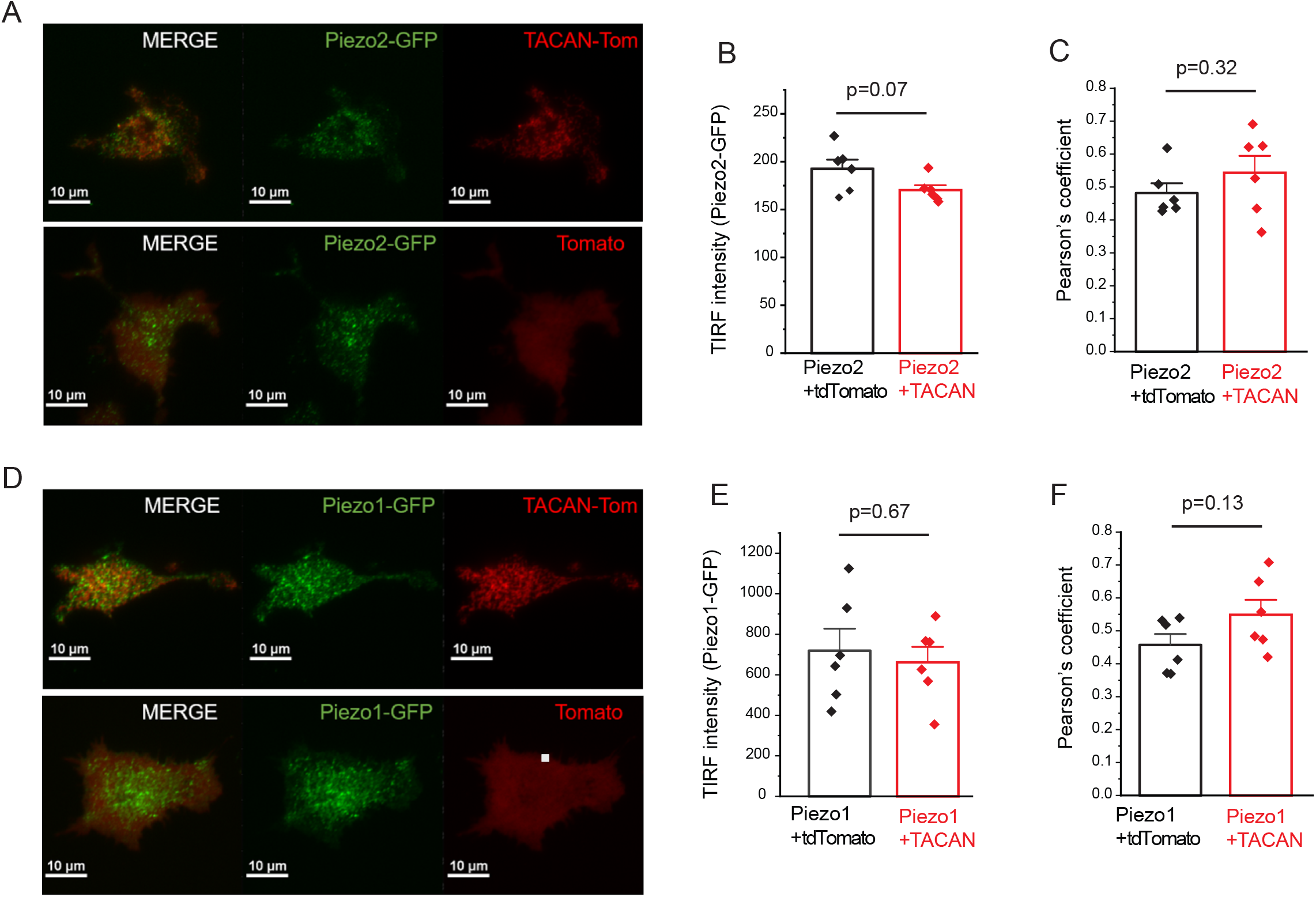
TACAN shows only weak colocalization with Piezo1 or Piezo2, and does not change their cell surface expression. TIRF microscopy was performed as described in the methods section. **(A)** Representative TIRF images for HEK293 cell transfected with tdTomato-TACAN and GFP-Piezo2 (top) and tdTomato and GFP-Piezo2 (Bottom). **(B)** Summary data for the fluorescence intensity for GFP-Piezo2 in the TIRF mode for cells cotransfected with tdTomato, or tdTomato-TACAN. **(C)** Pearson’s coefficient for colocalization of tdTomato-TACAN with GFP-Piezo2, and tdTomato with GFP-Piezo2. **(D)** Representative TIRF images for HEK293 cell transfected with tdTomato-TACAN and GFP-Piezo1 (top) and tdTomato and GFP-Piezo1 (Bottom). **(E)** Summary data for the fluorescence intensity for GFP-Piezo1 in the TIRF mode for cell cotransfected with tdTomato, or tdTomato-TACAN. **(F)** Pearson’s coefficient for colocalization of tdTomato-TACAN with GFP-Piezo1, and tdTomato with GFP-Piezo1. Bar graphs show mean ± SEM and scatter plots. Individual symbols show the average value of cells for one coverslip (5-22 cells / coverslip) from two independent transfection. Statistical significance was calculated with two sample t-test.

Both GFP-Piezo2 and GFP-Piezo1 showed punctate localization in the TIRF images (**Fig. 3A,D**.), in accordance with earlier reports (Ellefsen et al., 2019; Gottlieb and Sachs, 2012; Jiang et al., 2021; Ridone et al., 2020). tdTomato-TACAN also showed inhomogenous distribution, and displayed a weak colocalization with Piezo2 and Piezo1 with Person’s coefficients of ∼0.55. tdTomato alone showed a more homogenous distribution in TIRF images, and its Pearson’s coefficient with Piezo2 and Piezo1 was only slightly, and not statistically significantly lower than that for tdTomato-TACAN (**Fig3. C,F**).

Piezo2 has been shown to be modulated by both the actin cytoskeleton and microtubules (Chang and Gu, 2020; Jia et al., 2016). To test if TACAN acts via altering the organization of the plasma membrane associated cytoskeleton, we labeled HEK293 cells expressing tdTomato-TACAN and GFP-Piezo1 or GFP-Piezo2 with the cell permeable live cell dyes Sir-Actin and Spy650-tubulin and performed TIRF microscopy. Both actin **(Figure 3 Figure Supplement 1**) and tubulin **(Figure 3 Figure Supplement 2**) showed a very weak colocalization with Piezo2 and Piezo1 (Pearson’s coefficients between 0.1-0.4), and TACAN did not change this colocalization. TACAN also did not change the intensity of actin or tubulin labeling. These data indicate that TACAN does not act via altering the cytoskeletal elements adjacent to the plasma membrane.

Next, we tested if TACAN is expressed in the same neurons as Piezo2 using RNAscope fluorescence in situ hybridization. We performed triple labeling experiments with Piezo2, TACAN and one of the following neuronal markers: Neurofilament heavy chain (NFH), Tyrosine hydroxylase (TH), Transient receptor potential vanilloid 1 (TRPV1) and calcitonin gene-related peptide 2 (CGRP2) (**Fig. 4**). TACAN was expressed in >90% of neurons. Piezo2 was found in 60-70% of neurons and >95% of the Piezo2 positive neurons also expressed TACAN. Piezo2 showed a variable level of coexpression with the different neuronal markers, the highest being NFH, the lowest being TRPV1 (**Fig. 4 B,E,H,K**). We plotted the intensity of the Piezo2 signal as a function of the TACAN signal of each individual neuron that also showed staining with the neuronal markers (**Fig. 4 C,F,I,L**). Each neuronal subpopulation displayed cells with different TACAN/Piezo2 ratios, with a tendency of having cell populations with higher Piezo2 and lower TACAN, as well as higher TACAN and lower Piezo2 expression levels. These data suggest that TACAN/Piezo2 ratios may determine Piezo2 current levels.

**Figure 4.**
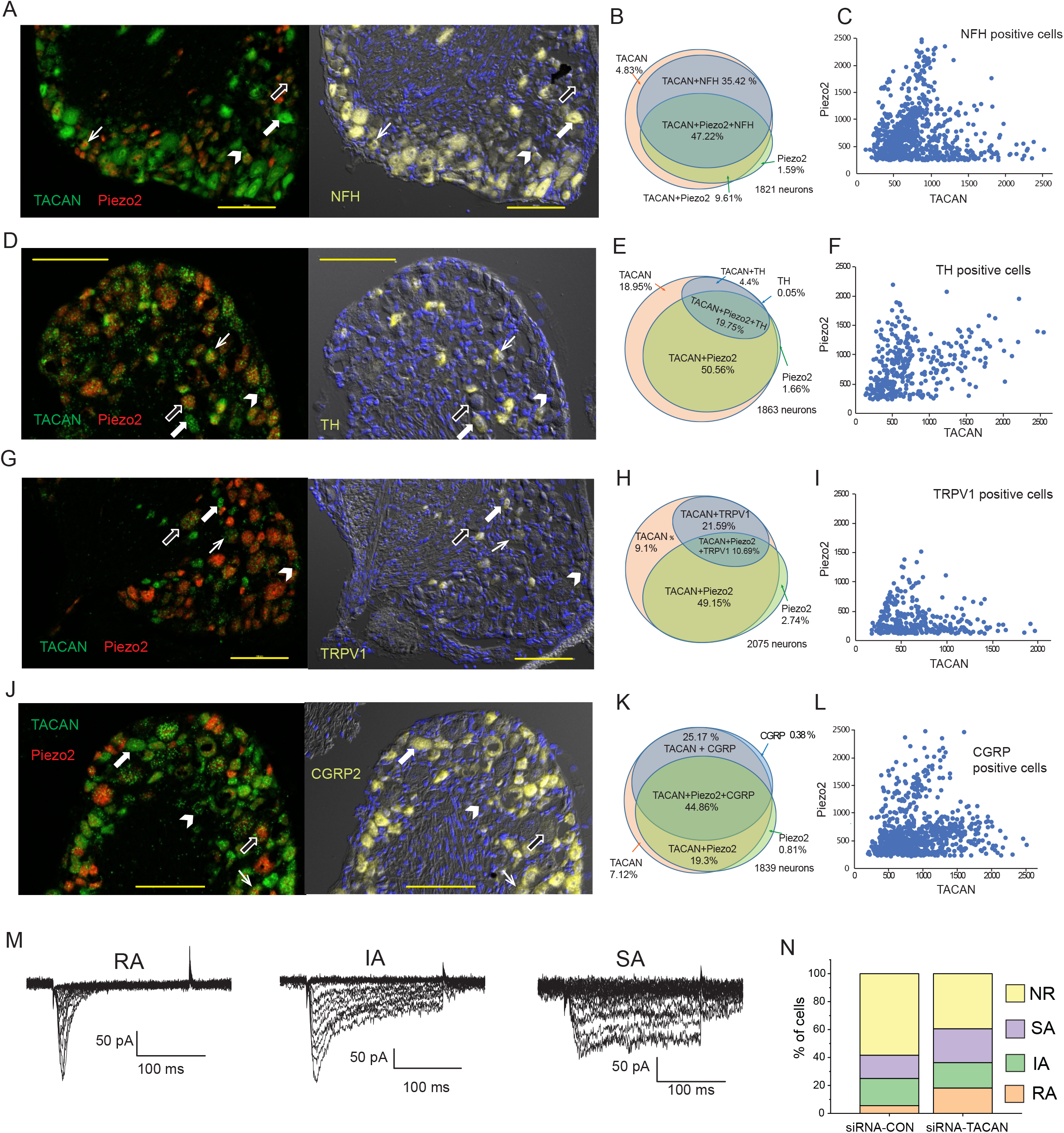
TACAN and Piezo2 are coexpressed in various DRG populations. RNAScope fluorescence in situ hybridization on mouse DRGs was performed as described in the methods section, with probes for TACAN, Piezo2 and four different neuronal markers. The data represent 3 independent DRG preparations, and 2-3 slices per condition for each preparation. Arrows in representative images show cells that express: TACAN and neuronal marker (white wide arrow), TACAN and Piezo2 and neuronal marker (white arrow), TACAN (arrow head) and TACAN+Piezo2 (black wide arrow). Horizontal yellow lines indicate 100 μm on each image. **(A)** Left panel: representative image for DRGs labeled with TACAN (green) and Piezo2 (red). Right panel, the same section labeled with Neurofilament heavy chain (NHF) (yellow). The section was also stained with DAPI to label nuclei. **(B)** Venn diagram showing coexpression of TACAN, Piezo2 and NHF. **(C)** Intensity of Piezo2 labeling as a function of TACAN signal for individual cells that were positive for NHF. For both TACAN and Piezo2, cells that were below the threshold for counting as positive for TACAN or Piezo2, are not shown. **(D)** Representative images for DRGs labeled with TACAN (green) and Piezo2 (red) and tyrosine hydroxylase (TH) (yellow). **(E)** Venn diagram showing coexpression of TACAN, Piezo2 and TH. **(F)** Intensity of Piezo2 labeling as a function of TACAN signal for individual cells that were positive for TH. **(G)** Representative images for DRGs labeled with TACAN (green) and Piezo2 (red) and TRPV1 (yellow). **(H)** Venn diagram showing coexpression of TACAN, Piezo2 and TRPV1. **(I)** Intensity of Piezo2 labeling as a function of TACAN signal for individual cells that were positive for TRPV1. **(J)** Representative images for DRGs labeled with TACAN (green) and Piezo2 (red) and CGRP2 (yellow). **(K)** Venn diagram showing coexpression of TACAN, Piezo2 and CGRP2. **(L)** Intensity of Piezo2 labeling as a function of TACAN signal for individual cells that were positive for CGRP2. **(M)** Representative traces for whole cell patch clamp recording in DRG neurons for Rapidly Adapting (RA), Intermediate Adapting (IA) and Slowly Adapting (SA) mechanically activated currents, evoked by mechanical indentation of the cells with 0.4 μm increments. **(N)** Percentage of cells displaying RA, IA and SA currents, as well as non-responding cells (NR) in neurons transfected with an siRNA against TACAN, or control siRNA. The electrophysiology data are from 9 independent DRG neuron preparations and transfections, n=36 for control siRNA, and n=33 for TACAN siRNA.

To assess the role of endogenous TACAN, we transfected isolated DRG neurons with fluorescently labeled siRNA against TACAN, or non-coding control siRNA. We patched cells that displayed red fluorescence, and mechanically stimulated them with a blunt glass probe. **Figure 4M** shows representative current traces for rapidly adapting, intermediate adapting and slowly adapting mechanically activated currents. TACAN was proposed to mediate slowly adapting currents (Beaulieu-Laroche et al., 2020). In our hands the proportion of slowly adapting currents in DRG neurons transfected with siRNA against TACAN, did not decrease compared to cells that were transfected with control siRNA **(Fig. 4N)**. This finding argues against TACAN mediating slowly adapting mechanically activated currents in DRG neurons. On the other hand, the percentage of cells displaying rapidly adapting currents increased, which is compatible with TACAN acting as a negative regulator of Piezo2 in DRG neurons.

## DISCUSSION

Here we report that TACAN inhibits the activity of mechanically activated Piezo2 channels. TACAN was proposed to function as an ion channel responsible for slowly adapting mechanically activated currents in DRG neurons. Overexpression of TACAN increased mechanically activated currents in response to negative pressure applied through the patch pipette in the cell attached configuration in several cell lines (Beaulieu-Laroche et al., 2020). These currents however were very small, only 1-2 pA on average, approximately doubling the amplitudes of background currents observed in nontransfected cells (Beaulieu-Laroche et al., 2020). In our hands, overexpression of TACAN in Piezo1 deficient N2A cells did not increase mechanically activated currents evoked by negative pressures over background levels in the cell attached configuration, but overexpression of Piezo1 with or without TACAN induced a 5.5 fold increase over background current levels. Overexpression of TACAN, on the other hand, evoked a robust decrease in Piezo2 currents evoked by indentation of the cell membrane with a blunt glass probe in the whole cell configuration.

In the whole cell configuration neither the original publication describing TACAN (Beaulieu-Laroche et al., 2020), nor we observed increased mechanically activated currents after overexpression of TACAN. Treatment of DRG neurons in the TRPV1 lineage with siRNA against TACAN on the other hand decreased the proportion of mechanically activated currents with ultra-slow inactivation kinetics in the whole cell configuration (Beaulieu-Laroche et al., 2020). In our hands, TACAN siRNA had no effect on the proportion of slowly adapting MA currents, but it increased the proportion of rapidly adapting MA currents in DRG neurons, which is consistent with TACAN suppressing Piezo2 activity. In a recent article single cell RNA sequencing experiments showed that TACAN showed the highest expression in DRG neurons that did not show responses to mechanical indentations, also raising doubts about TACAN serving as a mechanically activated ion channel in DRG neurons (Michel et al., 2020).

TACAN was shown by immunohistochemistry to be expressed predominantly in small non-peptidergic DRG neurons with a significant expression also in Tyrosine hydroxylase positive neurons (Beaulieu-Laroche et al., 2020). In our hands, TACAN expression with RNAScope in situ hybridization was detected in more than 90% of neurons, both in cells expressing NHF, a marker of myelinated neurons, and in neurons expressing CGRP2, TRPV1 or TH. TACAN and Piezo2 expression also showed substantial overlap, with varying TACAN/Piezo2 ratios, indicating that TACAN may, in principle, regulate endogenous Piezo2 channels in DRG neurons.

When TACAN was conditionally knocked out from non-peptidergic DRG neurons that express Mrgprd in a tamoxifen-inducible fashion, mice showed reduced nocifensive responses to von-Frey filaments (greater or equal to 1g), but they retained reflexive withdrawal (Beaulieu-Laroche et al., 2020). Paw withdrawal from painful pinprick stimuli on the other hand was not affected by TACAN deletion (Beaulieu-Laroche et al., 2020). TACAN knockdown by spinal intrathecal injection of antisense oligodeoxynucleotides into rats, reduced inflammatory, but not chemotherapy-induced mechanical hyperalgesia (Bonet et al., 2020). These data support the idea that TACAN plays a role in detection of some, but not all forms noxious mechanical stimuli.

Our data however indicate that TACAN inhibits Piezo2 currents. Can this be reconciled with these behavioral findings? As mentioned earlier, the role of Piezo2 in detecting noxious mechanical stimuli is complex. One study found that paradoxically, conditional deletion of Piezo2 in DRG neurons using an Advillin-cre mouse line reduced the threshold to painful mechanical stimuli in the Randall-Selitto test, even though the mice had defective gentle touch (Zhang et al., 2019). Ectopic expression of Piezo1 in DRG neurons on the other hand decreased sensitivity in the Randall-Selitto test (Zhang et al., 2019). These data indicate that activation of Piezo2 may inhibit mechanical pain. This is compatible with classical gate control theory of pain (Melzack and Wall, 1965), where stimulation of light touch receptors reduce pain. In this framework, increasing Piezo2-mediated currents by TACAN knockdown, can potentially reduce pain.

What is the mechanism of Piezo2 inhibition by TACAN? Our TIRF data indicate that TACAN co-expression does not decrease cell surface expression of Piezo2. While TIRF imaging does not have sufficient resolution to detect protein-protein interaction, the weak colocalization of TACAN with Piezo2 makes it unlikely that the mechanism is direct interaction with the channel. TACAN also did not induce a major reorganization of the actin and tubulin cytoskeleton, and it did not inhibit the activity of two other mechanosensitive channels Piezo1 and TREK1. These data indicate that the inhibition of Piezo2 is not due to a general change in mechanical properties of the cell, or a general decrease of the ability of the cell to transduce mechanical forces to ion channels. TACAN was also characterized earlier as a fat specific Nuclear Envelope Transmembrane protein 29 (NET29), and it was shown that its overexpression can alter gene expression (de Las Heras et al., 2017). Thus it is possible that TACAN increases or decreases the expression of other proteins that regulate Piezo2 properties. Recent preprints showed that the structure of TACAN determined by cryoEM showed similarity to the fatty acid elongase ELOVL, and the structure of TACAN also contained a coenzyme-A molecule (Niu et al., 2021; Rong et al., 2021; Xue et al., 2021). Piezo2 as well as Piezo1 were shown to be regulated by a variety of lipids (Borbiro et al., 2015; Narayanan et al., 2018; Romero et al., 2020; Romero et al., 2019), therefore it is possible that TACAN modulates Piezo2 activity through modifying the lipid content of the cell. Exploring this possibility will be a subject of future research.

**Figure 1 Figure Supplement 1.**
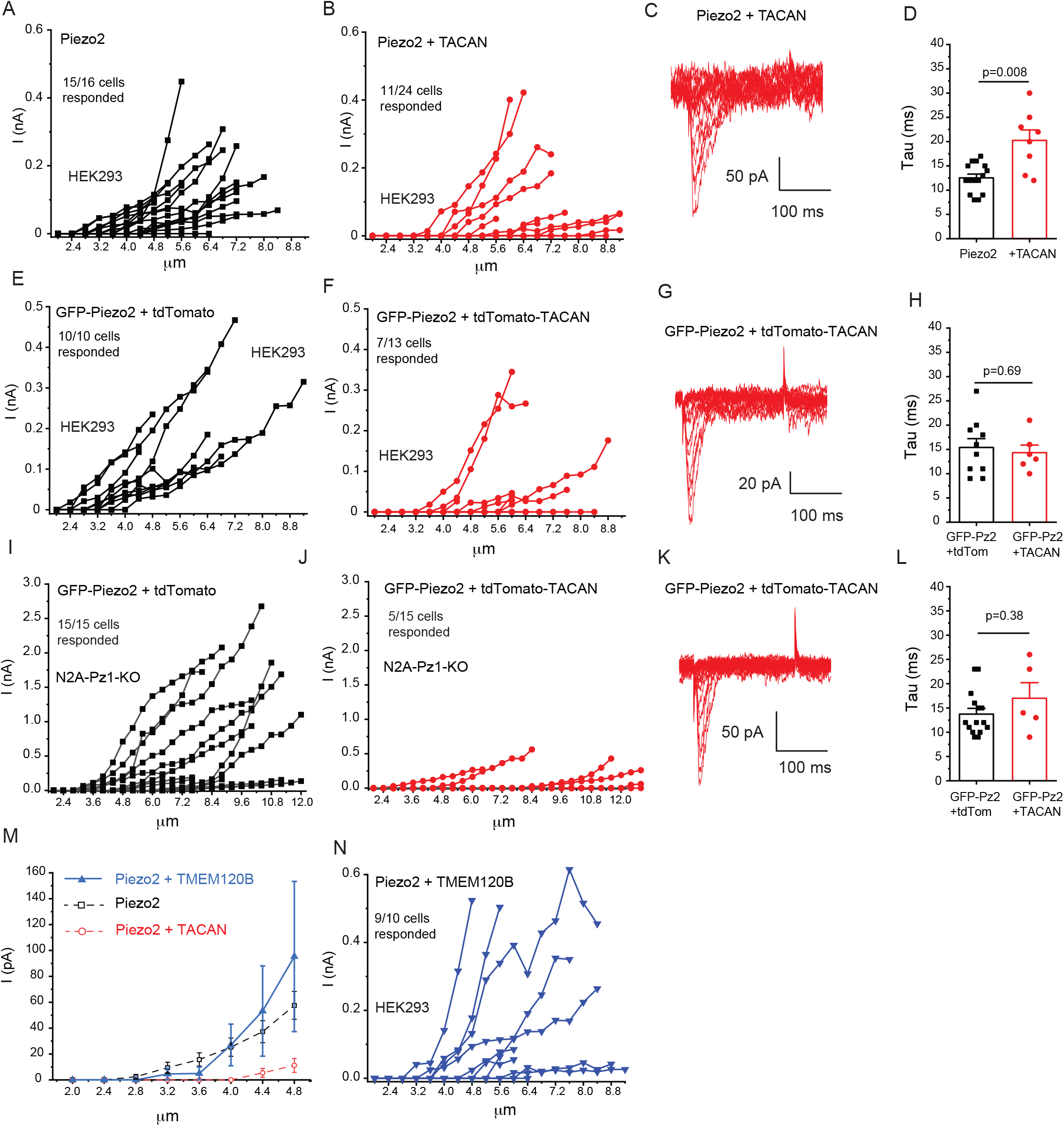
TACAN, but not TMEM120B inhibits Piezo2 currents. Data from Figure 1 showing the full range of measurements with indentation depth increased until the seal was lost. **(A**,**B)** Data from HEK293 cells transfected with Piezo2 and TACAN. **(C)** Representative traces for increasing indentation depth till 7.2 μm. **(D)** Inactivation time constant (Tau) for the same measurements. The inactivation kinetics from MA currents were measured by fitting the MA current with an exponential decay function, which measured the inactivation time constant (Tau). We used the currents evoked by the third stimulation after the threshold in most experiments, except in cells where only the two largest stimuli evoked a current, where we used the current evoked by the largest stimulus, provided it reached 40 pA. Statistical significance was calculated with two sample t-test. **(E**,**F)** Data from HEK293 cells transfected with GFP-Piezo2 and tdTomato-TACAN. **(G)** Representative traces for increasing indentation depth till 8.4 μm. **(H)** Inactivation time constant (Tau) for the same measurements. Statistical significance was calculated with two sample t-test. **(I**,**J)** Data from Piezo1 deficient N2A cells transfected with GFP-Piezo2 and tdTomato-TACAN. **(K)** Representative traces for increasing indentation depth till 11.6 μm. **(L)** Inactivation time constant (Tau) for the same measurements. Statistical significance was calculated with the Mann-Whitney-test. **(M)** Whole cell patch clamp data from HEK293 cells transfected with Piezo2 and TMEM120B, mean ± SEM of current amplitudes as a function of indentation depth are shown. For Piezo2 alone and Piezo2 + TACAN data were replotted from Figure 1 (dashed lines). **(N)** Data for full the range of indentations.

**Figure 2 Figure Supplement 1.**
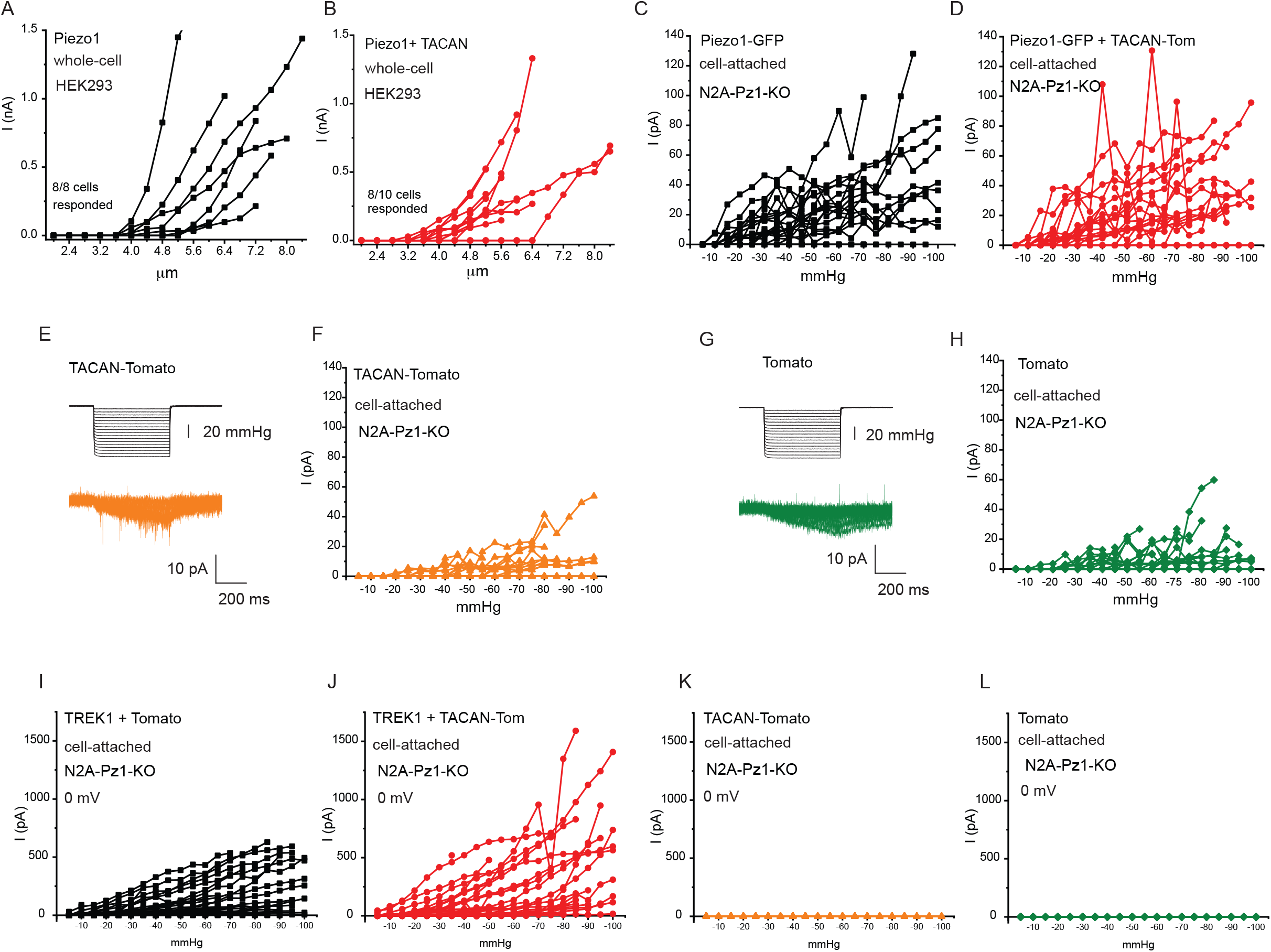
TACAN does not inhibit Piezo1 and TREK1 currents. Data from Figure 2 showing the full range of measurements with indentation depth or negative pressure increased until the seal was lost. **(A**,**B)** Data from whole cell patch clamp experiments in HEK293 cells transfected with Piezo1 alone or TACAN + Piezo1. **(C**,**D)** Data from cell attached patch clamp experiments in Piezo1 deficient N2A cells transfected with Piezo1 alone or TACAN + Piezo1. **(E)** Representative traces for cell-attached patch clamp experiments in N2A cell transfected with tdTomato-TACAN. **(F)** Full range of data from cell attached patch clamp experiments in N2A cell transfected with tdTomato-TACAN. **(G)** Representative traces for cell-attached patch clamp experiments in N2A cell transfected with tdTomato. **(H)** Full range of data from cell-attached patch clamp experiments in N2A cell transfected with tdTomato. **(I-L)** Full range of data from cell attached patch clamp experiments at 0 mV in N2A cells transfected with TREK1 + tdTomato (I), TREK1+tdTomato-TACAN (J), tdTomato-TACAN alone (K) and tdTomato alone.

**Figure 3 Figure Supplement 1.**
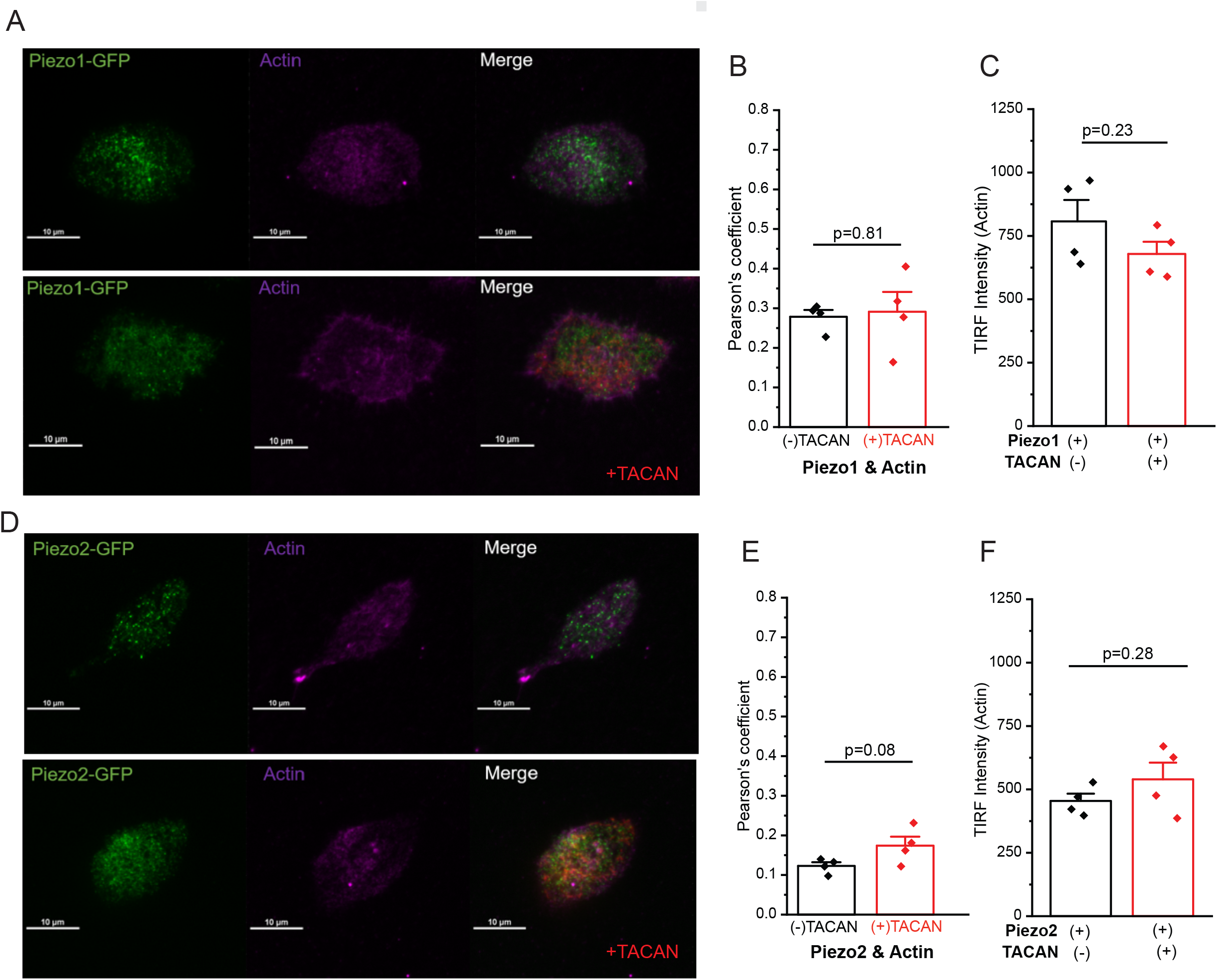
TACAN expression does not affect the actin cytoskeleton. HEK293 cells were transfected with tdTomato-TACAN and GFP-Piezo1, or GFP-Piezo2, labeled with Sir-Actin and TIRF images were obtained as described in the methods section. **(A)** Representative TIRF images for GFP-Piezo1 and Sir-Actin. **(B)** Pearson’s coefficient for colocalization of GFP-Piezo1 and Sir-Actin with and without TACAN. **(C)** TIRF intensity of Sir-Actin with and without TACAN. **(D)** Representative TIRF images for GFP-Piezo2 and Sir-Actin. **(E)** Pearson’s coefficient for colocalization of GFP-Piezo2 and Sir-Actin with and without TACAN. **(F)** TIRF intensity of Sir-Actin with and without TACAN.

**Figure 3 Figure Supplement 2.**
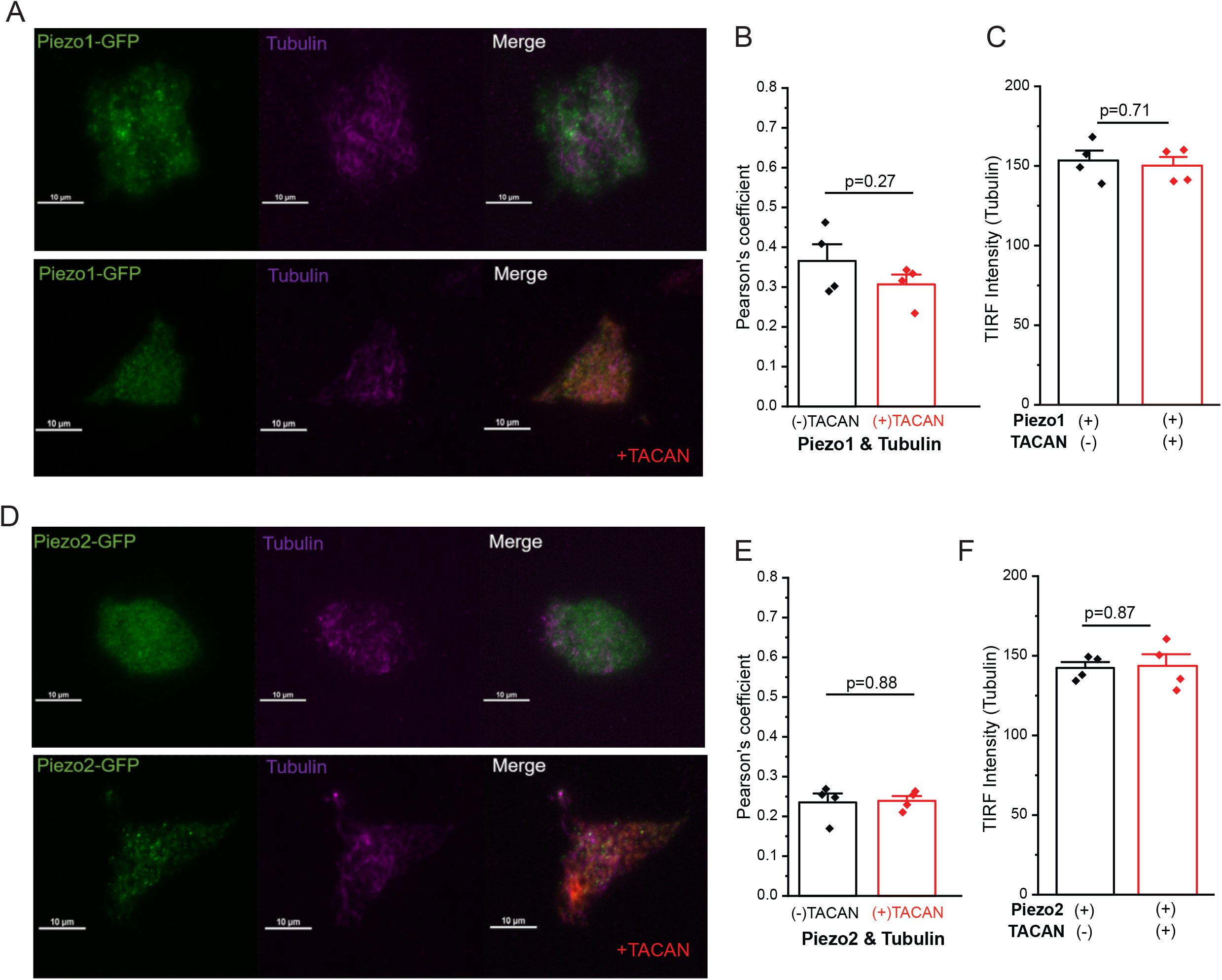
TACAN expression does not affect the tubulin cytoskeleton. HEK293 cells were transfected with tdTomato-TACAN and GFP-Piezo1 or GFP-Piezo2, labeled with Spy650-tubulin and TIRF images were obtained as described in the methods section. **(A)** Representative TIRF images for GFP-Piezo1 and Spy650-tubulin ctin. **(B)** Pearson’s coefficient for colocalization of GFP-Piezo1 and Spy650-tubulin with and without TACAN. **(C)** TIRF intensity of Spy650-tubulin with and without TACAN. **(D)** Representative TIRF images for GFP-Piezo2 and Spy650-tubulin. **(E)** Pearson’s coefficient for colocalization of GFP-Piezo2 and Spy650-tubulin with and without TACAN. **(F)** TIRF intensity of Spy650-tubulin with and without TACAN.

## MATERIALS AND METHODS

### KEY RESOURCES TABLE

**KEY RESOURCES TABLE:**
| **Reagent type** | **Designation** | **SOURCE** | **IDENTIFIER** | **Additional information** |
| --- | --- | --- | --- | --- |
| Chemical | NaCl | Fisher Scientific | S-271 |  |
| Chemical | KCl | MilliporeSigma | 60128-250G-F |  |
| Chemical | CaCl<sub>2</sub> • 2H<sub>2</sub>O | MilliporeSigma | C5080-500G |  |
| Chemical | MgCl<sub>2</sub> • 6H<sub>2</sub>O | MilliporeSigma | M2670-500G |  |
| Chemical | HEPES | MilliporeSigma | H3375-500G |  |
| Chemical | TEA-Cl • H<sub>2</sub>O | Acros Organics | AC150900250 |  |
| Chemical | K-gluconate | MilliporeSigma | G4500-500G |  |
| Chemical | EGTA | MilliporeSigma | E4378-250G |  |
| Chemical | glucose | J.T. Baker | 1916-05 |  |
| Chemical | Penicillin-streptomycin | Gemini | 400-109 |  |
| Chemical | Minimal Essential Medium | Gibco | 11095-080 |  |
| Chemical | Dulbecco's Modified Eagle's medium F-12 (HAM) | Gibco | 11320-033 |  |
| Chemical | Fetal Bovine Serum | Gemini | 100-500 |  |
| Chemical | Effectene | Qiagen | 301425 | Transfection reagent |
| Chemical | Sir-Actin | Cytoskeleton Inc | CY-SC001 | Labels F-actin in live cells |
| Chemical | Spy650-tubulin | Cytoskeleton Inc | CY-SC503 | Labels microtubules in live cells |
| Chemical | fluorescent siRNA TMEM120A | Qiagen | S101311919 |  |
| Chemical | AllStars Negative control siRNA | Qiagen | 1027294 |  |
| Chemical | HiPerFect | Qiagen | 301704 | Transfection reagent |
| Chemical | RNAscope® 2.5 LS Multiplex Reagent Kit | Advanced Cell Diagnostics | 323110 |  |
| Chemical | RNAscope® LS 2.5 Probe- Mm-TMEM120A | Advanced Cell Diagnostics | 513211 |  |
| Chemical | RNAscope® 2.5 LS Probe- Mm- Piezo2-C2 | Advanced Cell Diagnostics | 400191-C2 |  |
| Chemical | RNAscope® 2.5 LS Probe-Mm-NefH-C3 | Advanced Cell Diagnostics | 443671-C3 |  |
| Chemical | RNAscope® 2.5 LS Probe-Mm-TH-C3 | Advanced Cell Diagnostics | 317621-C3 |  |
| Chemical | RNAscope® 2.5 LS Probe-Mm-TRPV1-C3 | Advanced Cell Diagnostics | 313331-C3 |  |
| Chemical | RNAscope® 2.5 LS Probe-Mm-CalcB-C3 | Advanced Cell Diagnostics | 425511-C3 |  |
| Cell line (human) | HEK293 | ATCC | CRL-1573 |  |
| Cell line | Piezo1 deficient Neuro2A | (Moroni et al., 2018) | N/A | Kind gift from Drs. Valeria Vasquez and Gary Lewin |
| Organisms/Strains | C57Bl6 mice | Jackson Laboratories |  |  |
| Recombinant DNA reagent | Plasmid: Piezo2 pCDNA | (Coste et al., 2010) |  | Kind gift from Dr. Ardem Patapoutian |
| Recombinant DNA reagent | Plasmid: Piezo1 IRES GFP | (Coste et al., 2010) |  | Kind gift from Dr. Ardem Patapoutian |
| Recombinant DNA reagent | Plasmid: Tmem120A (TACAN) | Origene | MR205146 |  |
| Recombinant DNA reagent | Plasmid: Tmem120B | Origene | MR205067 |  |
| Recombinant DNA reagent | Plasmid : tdTomato-Tmem120A | Rutgers NJMS | N/A |  |
| Recombinant DNA reagent | Plasmid: GFP-Piezo2 | Rutgers NJMS | N/A |  |
| Recombinant DNA reagent | Plasmid: GFP-Piezo1 | Rutgers NJMS | N/A |  |
| Recombinant DNA reagent | Plasmid: TREK1 (KCNK2) | Origene | MR206535 |  |

### Cell Culture

Human Embryonic Kidney 293 (HEK293) cells were purchased from American Type Culture Collection (ATCC), Manassas, VA, (catalogue number CRL-1573), RRID:CVCL_0045. Cell identity was verified by STR analysis at ATCC. Additional cell authentication was not performed, but passage number of the cells was monitored, and cells were used up to passage number 25–30 from purchase, when a new batch of cells was thawed with low passage number. HEK293 cells were cultured in minimal essential medium (MEM) with 10% fetal bovine serum (FBS) and 100 IU/ml penicillin plus 100 µg/ml streptomycin in 5% CO_2_ at 37 °C. Cells were transfected with the Effectene reagent (Qiagen).

Piezo1 deficient N2A cells, in which endogenous Piezo1 was deleted by CRISPR (Moroni et al., 2018; Romero et al., 2020), were cultured in Dulbecco’s Modified Eagle Medium (DMEM), 5% penicillin–streptomycin, and 10 % fetal bovine serum (FBS). Cells were transfected with the Effectene reagent (Qiagen).

### Dorsal Root Ganglion neurons

DRG neurons were isolated as described previously (Borbiro et al., 2015; Del Rosario et al., 2020). All animal procedures were approved by the Institutional Animal Care and Use Committee at Rutgers NJMS. Mice were kept in a barrier facility under a 12/12 hr light dark cycle. Wild-type C57BL6 mice (2-to 4-month old) from either sex (Jackson laboratories) were anesthetized with i.p. injection of ketamine (100 mg/kg) and xylazine (10 mg/kg) and perfused via the left ventricle with ice-cold Hank’s buffered salt solution (HBSS; Life Technologies). DRGs were harvested from all spinal segments after laminectomy and removal of the spinal column and maintained in ice-cold HBSS for the duration of the isolation. Isolated ganglia were cleaned from excess nerve tissue and incubated with type 1 collagenase (2 mg/ml; Worthington) and dispase (5 mg/ml; Sigma) in HBSS at 37°C for 30 min, followed by mechanical trituration. Digestives enzymes were then removed after centrifugation of the cells at 100 g for 5 min.

Isolated DRG neurons were resuspended and seeded onto laminin coated glass coverslips; 2-3 hours after seeding, DRG neurons were transfected in serum free DMEM solution with predesigned fluorescently labelled siRNA directly against mouse TMEM120A (TACAN) from Qiagen-FlexiTube (cat. S101311919) or AllStars Neg. siRNA (cat. 1027294) at a final concentration of 200nM, using the HiPerFect transfection reagent from Qiagen. 24 hr after transfection, cells were incubated with fresh serum-containing media and incubated for an additional 40-48hr prior to whole cell patch electrophysiology. Transfected cells were identified by their fluorescence signal and selected as candidates for patch clamp experiments.

### Whole cell patch clamp

Whole-cell patch clamp recordings were performed at room temperature (22° to 24°C) as described previously (Borbiro et al., 2015). Briefly, patch pipettes were prepared from borosilicate glass capillaries (Sutter Instrument) using a P-97 pipette puller (Sutter instrument) and had a resistance of 2-7 MΩ. After forming gigaohm-resistance seals, the whole cell configuration was established, and the MA currents were measured at a holding voltage of -60 mV using an Axopatch 200B amplifier (Molecular Devices) and pClamp 10. Currents were filtered at 2 kHz using low-pass Bessel filter of the amplifier and digitized using a Digidata 1440 unit (Molecular Devices). All measurements were performed with extracellular (EC) solution containing 137 mM NaCl, 5 mM KCl, 1 mM MgCl_2_, 2 mM CaCl_2_, 10 mM HEPES and 10 mM glucose (pH adjusted to 7.4 with NaOH). The patch pipette solution contained 140 mM K^+^ gluconate, 1 mM MgCl_2_, 2 mM Na_2_ATP, 5 mM EGTA and 10 mM HEPES (pH adjusted to 7.2 with KOH).

Mechanically activated currents were measured from isolated DRG neurons, transiently transfected HEK293 cells and transiently transfected Piezo1 deficient N2A cells, as previously described (Borbiro et al., 2015). Briefly, Mechanical stimulation was performed using a heat-polished glass pipette (tip diameter, about 3 µm), controlled by a piezo-electric crystal drive (Physik Instrumente) positioned at 60° to the surface of the cover glass. The probe was positioned so that 10-µm movement did not visibly contact the cell but an 11.5-µm stimulus produced an observable membrane deflection.

Measurements from cells that showed significant swelling after repetitive mechanical stimulation, or had substantially increased leak current were discarded. For the DRG neuron experiments, to categorize mechanically activated currents as rapidly, intermediate and slowly adapting, we used the criteria from our previous publication (Borbiro et al., 2015). We considered currents to be rapidly adapting, if they fully inactivated before end of the 200 ms mechanical stimulus. The inactivation time constants (tau) of these currents were 17.95 ± 2.4 ms, similar to that in our earlier work (Borbiro et al., 2015; Del Rosario et al., 2020). Intermediate adapting currents did not fully inactivate, but the leftover current at the end of the mechanical simulation was less that 50% of the peak current. The inactivation time constant for these currents were 56.38 ± 5.07 ms. Slow adapting currents also did not fully inactivate, and the leftover current at the end of the mechanical simulation was more that 50% of the peak current, and/or the time constant was over 100 ms. Many currents in this category could not be fit with an exponential decay function.

### Cell attached Patch clamp

Whole-cell patch clamp recordings were performed at room temperature (22° to 24°C) similarly to described previously (Lewis and Grandl, 2015). After forming gigaohm-resistance seals MA currents were measured at a holding voltage of -80 mV for Piezo1 currents and at 0 mV for TREK1 currents, using an Axopatch 200B amplifier (Molecular Devices) and pClamp 10. Currents were filtered at 2 kHz using low-pass Bessel filter of the amplifier and digitized using a Digidata 1440 unit (Molecular Devices). Mechanical stimulation in cell attached patches was performed using a high-speed pressure clamp (Besch et al., 2002) (HSPC-1, ALA Scientific) controlled by pClamp 11.1 software (Molecular Devices) as described earlier (Borbiro et al., 2015).

All measurements were performed with a bath solution containing, 140 mM KCl, 1 mM MgCl_2_, 10 mM HEPES and 10 mM glucose (pH adjusted to 7.4 with KOH) to bring the membrane potential of the cells close to zero. The patch pipette solution contained 130 mM NaCl, 5 mM KCl, 1 mM MgCl_2_, 1 mM CaCl_2_, 10 mM HEPES and 10 mM TEA-Cl (pH adjusted to 7.4 with NaOH).

### Total Internal Reflection Fluorescence (TIRF) Microscopy

HEK293 cells were transiently transfected with cDNA encoding tdTomato-TACAN or tdTomato, and GFP-Pieoz1 or GFP-Piezo2. The next day, the transfected cells were plated on poly-L-lysine-coated 35-mm round coverslip (#1.5 thickness) (Fisher Scientific, Waltham, MA). The cells were used for TIRF imaging 2 days after transfection. Cells plated on the coverslip were placed into a recording chamber filled with extracellular solution containing (in mM) 137 NaCl, 5 KCl, 1 MgCl_2_, 10 HEPES and 10 glucose (pH 7.4). TIRF images were obtained at room temperature using a Nikon Eclipse Ti2 microscope (Tokyo, Japan). Fluorescence excitation was performed using a 15-mW solid state 488 nm and 561 nm laser at 90% of the maximal power through a CFI Apochromat TIRF 60X oil objective (NA of 1.49). Images were captured using an ORCA-Fusion Digital CMOS camera (Hamamatsu, Hamamatsu City, Japan) through emission filters 525/50nm, and 600/50nm for the green and red channel, respectively. The images were analyzed using Nikon NIS-Elements AR Analysis software and Image J.

To visualize actin and tubulin cytoskeleton, the cells were labeled with either Sir-Actin (CY-SC001), or Spy650-tubulin (CY-SC503) (Cytoskeleton Inc) according to the manufacturers instructions. Live cells were incubated for 1 hour at 37°C with either Sir-Actin (2 μM), or Spy650-tubulin (500x dilution of the stock solution recommended by the manufacturer). The cells were washed with the extracellular solution before imaging to remove the excess dyes. To visualize the probes the cells were illuminated by the 640 nm 15 mW solid state laser, and emission was collected using a 700/75nm emission filter.

### RNAscope *in situ* hybridization

DRG ganglia isolation was performed as described before with transcardial perfusion under deep anesthesia first with 10 ml of HBSS and after with 10 ml of 4% formaldehyde, L3-L5 DRGs were collected from mice; the tissues were post-fixed for 1 hour in 4 % formaldehyde and dehydrated in a series of gradient sucrose concentration (10, 20 and 30 %). After freezing in O.C.T compound block (Sakura Finetech) DRGs were sectioned into 12 µm thick slices, RNAscope assay was then carried out as previously described (Su et al., 2020).

Simultaneous detection of mouse RNA transcripts for *Piezo2, TACAN*, and various neuronal markers was performed on fixed, frozen DRG sections using Advanced Cell Diagnostics (ACD) RNAscope® Multiplex Fluorescent Reagent Kit v2 (cat number: 323110) according to the manufacturer’s instructions, and commercially available probes for Mm-TMEM120a (Cat number 513211), Mm-Piezo2 (cat number: 400191-C2), Mm-Trpv1 (cat number: 313331-C3), Mm-Th (cat number: 317621-C3), Mm-NefH (cat number: 443671-C3) and Calcb/CGRP2 (cat number: 425511-C3) were purchased from Advanced Cell Diagnostics.

In short, the sections were post-fixed in pre-chilled 4% PFA for 15 min at 4°C, washed 3 times with PBS for 5 min each before dehydration through 50%, 70 & 100% and 100% Ethanol for 5 min each. We then treated slides with a protease 4 for 20 min and washed in distilled water. Probe hybridization and signal amplification was performed according to manufacturer’s instructions. The following TSA plus fluorophores were used to detect corresponding RNAscope probes: Opal 520, 570 and 690 reagent kits (Akoya Biosciences). Cells were stained with DAPI (ACD) and mounted on the slide with Gold Antifade Mountant. Slides were imaged on a Nikon A1R confocal microscope with a 20x Plan Apo air objective, NA 0.75; and images were quantified in Nikon NIS-Elements. Totally 3 mouse were used in the experiment, from each animals were harvested 6 L3-5 DRG ganglions, after sectioning of the tissue 6 slices from each animals were treated and 2-3 slices from each animals were randomly selected for analysis, a total number of 2075 neurons for V1 probe, 1863 neurons for the TH probe, 1821 neurons for NF and 1839 neurons for CGRP2 probe were analyzed.

Neuronal cells borders were determined and segmented manually dependently on the level of the fluorescent probes signal and DIC images. Cells were considered as signal positive if the mean fluorescence signal in the ROI exceeded 0.5 times the standard deviation of the fluorescence signal in this channel.

### Statistics

Data are represented as mean ± standard error (S.E.M.) plus scatter plots. For normally distributed data statistical significance was calculated either with two sample t-test (two tailed), or analysis of variance (ANOVA), with Bonferroni post hoc test. Normality was assessed by the Shapiro Wilk test or the Lilliefors test. For non-normally distributed data Wilcoxon, or Mann-Whitney tests were used as appropriate. The specific tests for each experiment are described in the figure legends. Most statistical calculations and data plotting were performed using the Origin 2021 software.

## ACKNOWLEDGEMENTS

This study was supported by NIH grants NS055159 to T.R, and F31NS100484 and F99NS113422 to J.S.D.R. The Piezo1 IRES GFP and the Piezo2 PCDNA clones were kind gifts from Dr. Ardem Patapoutian, Scripps Research. The Piezo1 deficient Neuro2A cell line was a kind gift from Dr. Valeria Vasquez (University of Tennessee, Health Science Center) with permission from Dr. Gary Lewin (Max Delbruck Center for Molecular Medicine).

## Data Availability

All data generated or analyzed during this study are included in the manuscript and supporting files.

